# Evolutionary quantitative genomics of *Populus trichocarpa*

**DOI:** 10.1101/026021

**Authors:** Ilga Porth, Jaroslav Klápště, Athena D. McKown, Jonathan La Mantia, Robert D. Guy, Pär K. Ingvarsson, Richard Hamelin, Shawn D. Mansfield, Jüergen Ehlting, Carl J. Douglas, Yousry A. El-Kassaby

## Abstract

Forest trees generally show high levels of local adaptation and efforts focusing on understanding adaptation to climate will be crucial for species survival and management.

Merging quantitative genetics and population genomics, we studied the molecular basis of climate adaptation in 433 *Populus trichocarpa* (black cottonwood) genotypes originating across western North America. Variation in 74 field-assessed traits (growth, ecophysiology, phenology, leaf stomata, wood, and disease resistance) was investigated for signatures of selection (comparing *Q*_*ST*_-*F*_*ST*_) using clustering of individuals by climate of origin. 29,354 SNPs were investigated employing three different outlier detection methods.

*Narrow-sense Q*_*ST*_ for 53% of distinct field *Q*_*ST*_ traits was significantly divergent from expectations of neutrality (indicating *adaptive* trait variation); 2,855 SNPs showed signals of diversifying selection and of these, 118 SNPs (within 81 genes) were associated with adaptive traits (based on significant *Q*_*ST*_). Many SNPs were putatively pleiotropic for functionally uncorrelated adaptive traits, such as autumn phenology, height, and disease resistance.

Evolutionary quantitative genomics in *P. trichocarpa* provides an enhanced understanding regarding the molecular basis of climate-driven selection in forest trees. We highlight that important loci underlying *adaptive* trait variation also show relationship to climate of origin.

**Author summary:** Comparisons between population differentiation on the basis of quantitative traits and neutral genetic markers inform about the importance of natural selection, genetic drift and gene flow for local adaptation of populations. Here, we address fundamental questions regarding the molecular basis of adaptation in undomesticated forest tree populations to past climatic environments by employing an integrative quantitative genetics and landscape genomics approach. Marker-inferred relatedness was estimated to obtain the *narrow-sense* estimate of population differentiation in wild populations. We analyzed an unstructured population of common garden grown *Populus trichocarpa* individuals to uncover different extents of variation for a suite of field traits, wood quality and pathogen resistance with temperature and precipitation. We consider our approach the most comprehensive, as it uncovers the molecular mechanisms of adaptation using multiple methods and tests. We provide a detailed outline of the required analyses for studying adaptation to the environment in a population genomics context to better understand the species’ potential adaptive capacity to future climatic scenarios.

## Introduction

Knowledge about the genetic basis of *adaptive* quantitative traits in forest trees and genetic differentiation in response to selection facilitates the prediction of long-term responses to climate, but the genetic basis of adaptation is not comprehensively understood [1]. High levels of local adaptation due to consistent natural selection in a given environment resulted in local populations that have their highest fitness at their original provenance, and consequently, are differentiated from non-local populations. Within population diversity is fundamental to species survival in unpredictable environments, and therefore also relevant for conservation and forest management ([2]; [3]). Recent studies within forest trees have investigated the association of local climate and geography with either randomly identified loci (*Pinus taeda*: [4]; *Cryptomeria japonica*: [5], or candidate functional genes (*Picea abies*: bud set candidate genes, [6]; *Populus balsamifera*: flowering time candidate genes, [7]) to uncover genes underlying local adaptation. The genetic architecture underlying adaptive phenotypes of forest trees is generally highly complex (*e.g.* [8]). Therefore, untangling the relationships between adaptive loci and the role of climate in selection vs. neutral evolutionary processes is inherently difficult.

Evidence for potential adaptive significance of a genetic marker is often interpreted from ‘*F*_*ST*_ outlier’ analyses where genetic loci significantly differ in their allelic frequencies among populations. These ‘outliers’ can be efficiently detected using multilocus scans comparing patterns of nucleotide diversity and genetic differentiation to the simulated genome-wide neutral genetic background ([9]; [10]). For instance, this methodology has led to the detection of SNPs implicated in local climate adaptation in *Picea* ([11]; [12]; [13]). In order to obtain a detailed understanding of how populations have diverged in response to climate variation, such *F*_*ST*_ outliers can be tested for associations with an adaptive trait and an environmental variable to substantiate the evidence for their involvement in local adaptation ([14]; [15]). Integrating quantitative and population genomics is therefore essential to determine the degree to which genetic and phenotypic variation are driven by selection as opposed to neutral processes (*e.g.* genetic drift). Specifically, this allows for comprehensive information from genome-wide association studies (GWAS), *Q*_*ST*_ quantitative genetics analysis (*i*.*e*. ‘top-down’ approaches, [16]) and landscape population *F*_*ST*_ outlier analysis (*i*.*e*. ‘bottom-up’ approaches, [17]) be merged.

The existence of interaction effects among different loci within co-adapted gene complexes has long been recognized [18]. Yeaman (2013) suggested that ecological selection might even promote the physical clustering of locally adaptive loci through genomic rearrangements [19]. Landscape population genomics can identify genome regions significantly associated with spatial and temporal environmental gradients [3]. For instance, the study using natural *Arabidopsis* genotypes spanning the species’ range revealed that local adaptation might be maintained by independent target loci enriched for molecular processes that exhibit their major genetic effects within distinct local environments but are neutral in others [20]. The geographic variation in the degree to which a genetic region under selection responds is termed “conditional neutrality” [21] and suggests a given species has not uniformly responded to an environmental pressure or that the pressure is not equally active across a species range. Importantly, the assessment of local adaptation in this work on *Arabidopsis* involves the study of fitness traits such as fecundity and survival (viability) ([20]; [22]). In addition, there also exist traits that increase fitness in one environment, but reduce it in another. Ecological genetics can more easily explore the genetic changes over time in annuals (due to their short generation times) involving multiple generations studied under a changing environment ([23]; [15]). This is less feasible for long-lived forest trees. However, the estimation of quantitative genetic parameters using SNP marker-inferred relatedness estimation to obtain *narrow-sense* estimates of heritability and *Q*_*ST*_ in wild populations [24] can allow monitoring adaptive genetic responses along an ecological time-scale [15].

In this study, we integrated an extensive body of results on the genetics of wild *Populus trichocarpa* Torr. & A. Gray (black cottonwood) to understand adaptation to climate. All poplars, aspens, and cottonwoods (genus *Populus*) play important roles in natural ecosystems as pioneer species ([25]; [26]) and are economically important for various industrial products with an increasing role as bioenergy crops ([27]; [28]; [29]; [30]). *Populus* species are still largely undomesticated with very low population differentiation indicative of extensive long-distance intraspecific gene flow [31]. In western North America, *P. trichocarpa* has an extensive cordilleran range (31-62°N), yet with no clear north-south differentiation in genetic diversity (and no decreasing genetic diversity with latitude), consistent with the species’ colonization history from multiple potential glacial refugia [32]. Several studies have indicated subtle substructure in *P. trichocarpa* ([33]; [34]; [35]) relating to isolation-by-distance (IBD; *i*.*e*. the decrease of genetic similarity among populations with increasing geographical distance between these populations reflected in *continuous* patterns of genetic differentiation and allele frequency variation in the species [34] as opposed to natural barriers causing discrete local genetic clusters), introgression and adaptation [36]. We explored the extensive body of data on the genetics of *P. trichocarpa*, including genome-wide coverage of SNPs [35], and comprehensive GWAS results from wood characteristics [37], leaf rust fungus (*Melampsora* x*columbiana*) resistance [38], biomass, ecophysiology, leaf stomata and phenology traits [39]. We studied the divergence patterns of phenotypic variation and SNPs among distinct climate clusters in 433 unrelated *P. trichocarpa* genotypes originally collected throughout the northern two-thirds of the species’ latitudinal range (excluding the highly diverged Californian population Tahoe: [34], [40]). We tested whether phenotypic variation in traits was diverged among the climatic regions (based on non-neutral *Q*_*ST*_), as would be expected of adaptive variation. We then predicted that SNPs that are most diverged among different climatic regions would be associated with mapped genes that underlie *adaptive* phenotypic variation [13].

In brief, we used an integrative analysis of quantitative traits and genetic markers to investigate climate adaptation in wild *P. trichocarpa* populations, we developed an integrative approach through merging genomic-based datasets and results. (1) The effects of individual loci were first separated from confounding population effects using spatial PCA (sPCA) to investigate the presence of local and global genetic structures. Following this assessment of population structure using genetic markers showing evidence of only one single genetic structure, distinct population clusters were generated based on climatic factors and this subpopulation clustering was used in subsequent analyses (Fig. 1). (2) The genetic differentiation in quantitative traits (narrow-sense *Q*_*ST*_) among populations defined by climate clusters was calculated involving the estimation of relatedness based on genetic markers. (3) In parallel, the divergence of genetic markers (*F*_*ST*_ outlier analysis) among populations defined by climate clusters was assessed. (4) The significance of quantitative trait divergence among populations, as defined by climate clusters, was assessed by comparing the observed *Q*_*ST*_ values with the simulated distribution of *Q*_*ST*_-*F*_*ST*_ for a neutral trait. If the null hypothesis was rejected, the trait was considered adaptive. (5) GWAS results identifying the SNP variants underlying adaptive traits were incorporated. If these SNP variants also corresponded to loci under selection (employing four different outlier detection methods), then, the SNP variants were considered adaptive. This comprehensive analysis of genomic and phenotypic information underscores the necessity of merging multiple datasets to more fully understand evolutionary genomics of *P. trichocarpa*.

**Fig. 1.**
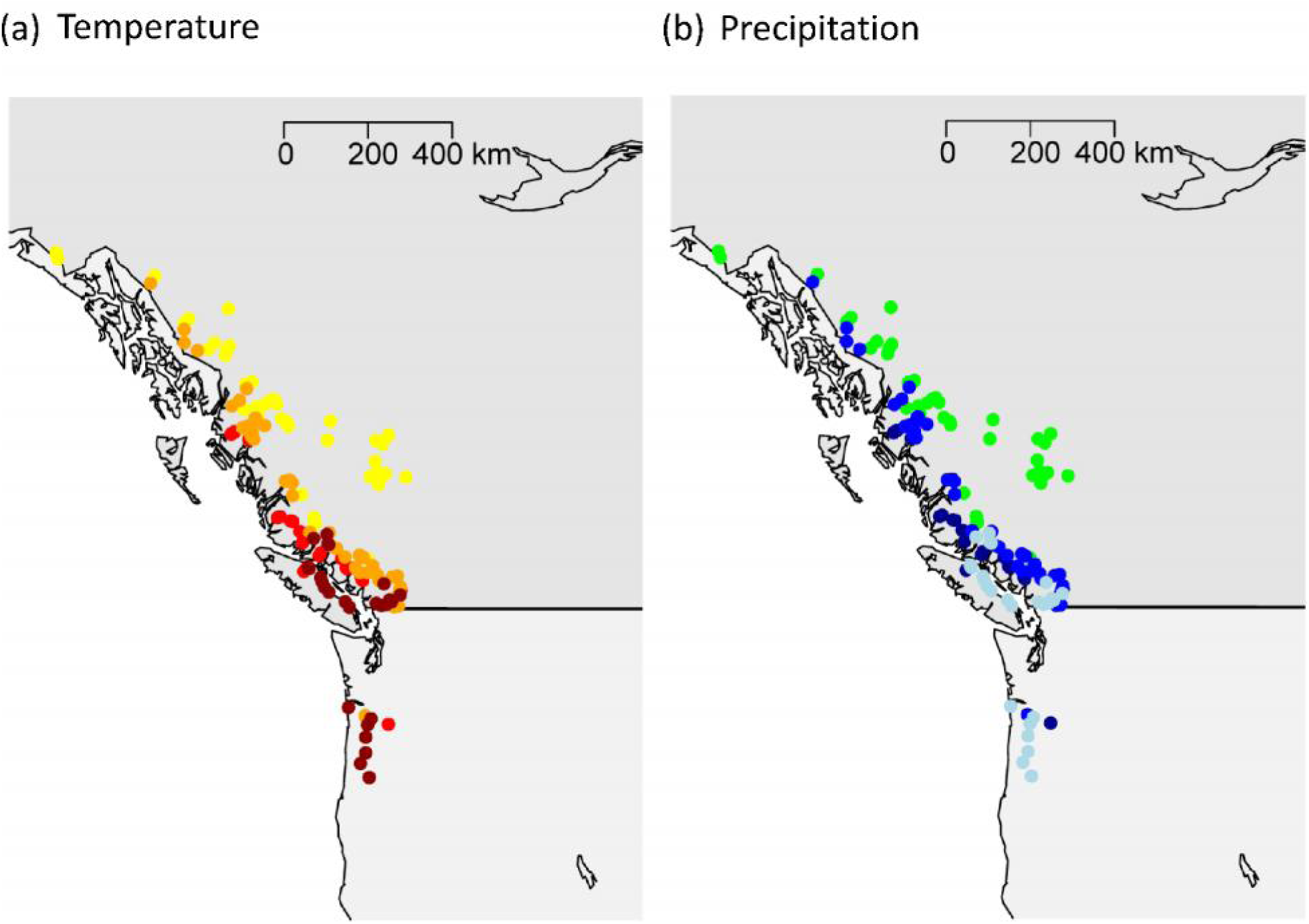
Geographical origins of 433 *P. trichocarpa* genotypes collected across 140 unique locations within the Pacific Northwest (British Columbia, Canada; Oregon, USA) and grouped into four distinct climate clusters using local temperature and precipitation records for location of origin. The climate regions were identified based on K-medoids clustering using the mean annual temperature (°C) between yrs 1971-2002 (MAT_1971-2002), the number of frost-free days (NFFD_1971-2002), and the mean annual precipitation (mm), observed between yrs 1971-2002 (MAP_1971-2002). Color coding is as follows: (a) population averages for MAT_1971-2002; NFFD_1971_2002: dark red (9.5°C; 287.1d); red (8.1°C; 267.2d); orange (6.4°C; 215.2d); yellow (4.2°C; 175.4d); (b) population average for MAP_1971-2002: dark blue (2805.9mm); blue (1571.8mm); light blue (1517.0mm); green (744.2mm). We note here that canonical correlations between geography and ecology were high (r=0.9 for the first canonical variable component). (TIFF)

## Results

### Population structure assessment

Negative eigenvalues from sPCA were negligible (Fig. 2), suggesting no local genetic clusters. By comparison, the presence of IBD was verified by large positive eigenvalues (Fig. 2). These results were further confirmed by the local and global tests within the “adegenet” program (see Methods). While, again, we did not detect local genetic structure in *P. trichocarpa* (local test *P*=0.937), we did identify global genetic structure attributed to IBD (global test *P*=0.001) that was observed across the entire population involving the 140 unique geographical locations represented by one randomly chosen genotype.

**Fig. 2.**
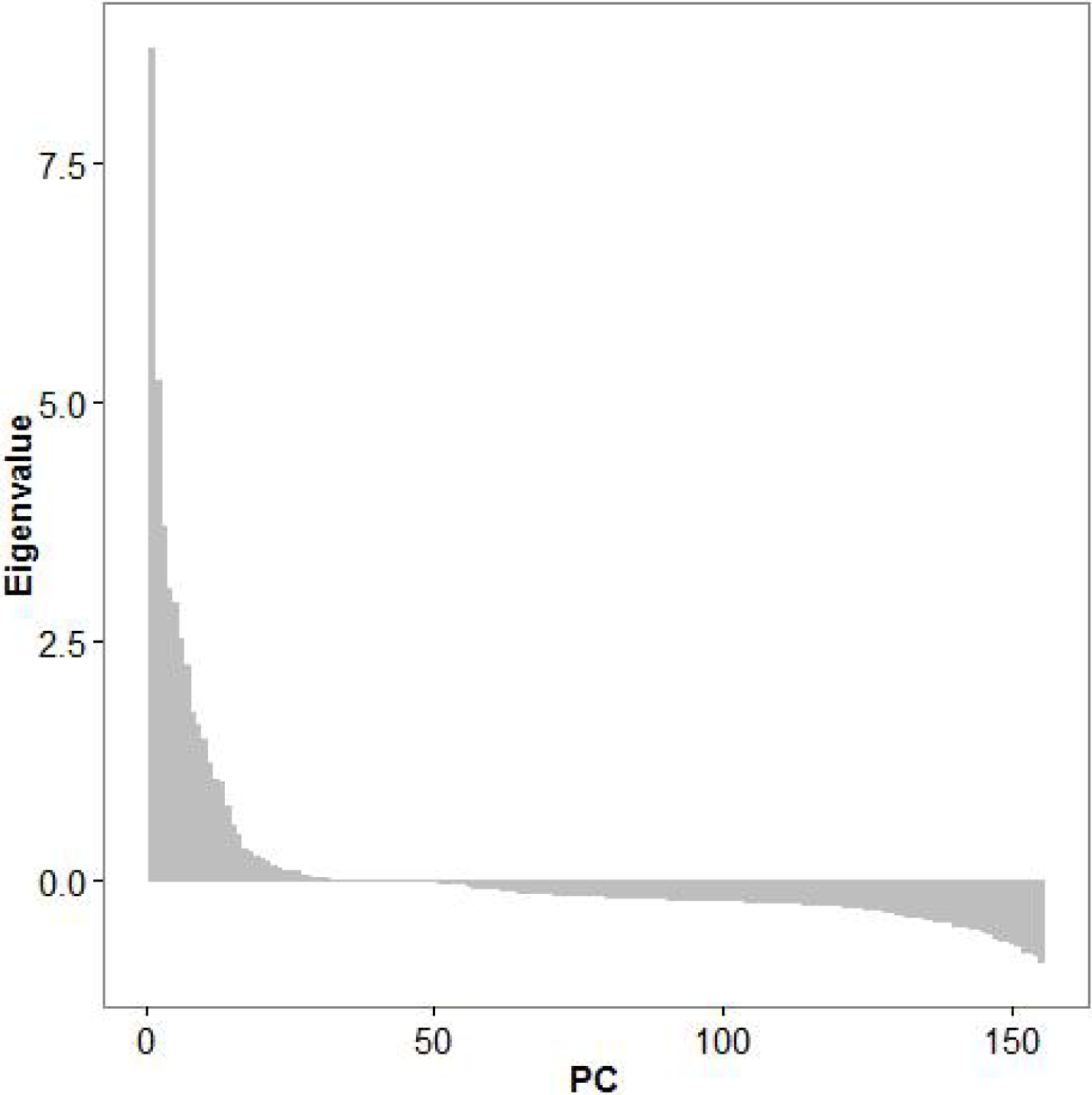
Identification of isolation-by-distance (IBD) among 433 *P. trichocarpa* genotypes based on spatial PCA. Large positive eigenvalues were indicative of IBD. (TIFF)

### Divergence of quantitative characters (*Q*_*ST*_) among climate clusters

We calculated *narrow-sense Q*_*ST*_ values for 74 distinct field-assessed traits for the study population. Assessments included 16 wood, 12 biomass, 14 phenology, 18 ecophysiological, 13 leaf stomata, and one rust resistance phenotype (Table S1). Observed *Q*_*ST*_ values for each trait were compared to the simulated distribution of *Q*_*ST*_-*F*_*ST*_ values for a neutral trait (simulating a range of possible demographic scenarios, see Methods). Among all traits, 53% (39/74 traits) had *Q*_*ST*_ values significantly different from zero and therefore were classified as *adaptive* (Table 1). The highest number of significant *Q*_*ST*_ values was observed among biomass traits (76%), phenology traits (70%), ecophysiology traits (64 %) and leaf rust resistance (100%). By comparison, only 25% of wood-based traits had significant *Q*_*ST*_ values. *Q*_*ST*_ values for traits that significantly diverged among the four climate clusters ranged from 0.03 (δ15N, *i*.*e*. stable nitrogen isotope ratio) to 0.26 (bole biomass). Among all tested traits, the climatic clusters best explained the phenotypic variation in phenology based on the *P*_*ST*_ values, ranging from 17% (100% leaf yellowing) to 24% (bud set). Among wood characteristics, two cell wall sugar traits (% galactose and % arabinose in dry wood) and two wood ultrastructure attributes (fiber length and microfibril angle) showed significant *Q*_*ST*_ values. The climatic clusters explained 13 and 12% of the arabinose and galactose content, respectively.

**Table 1.**
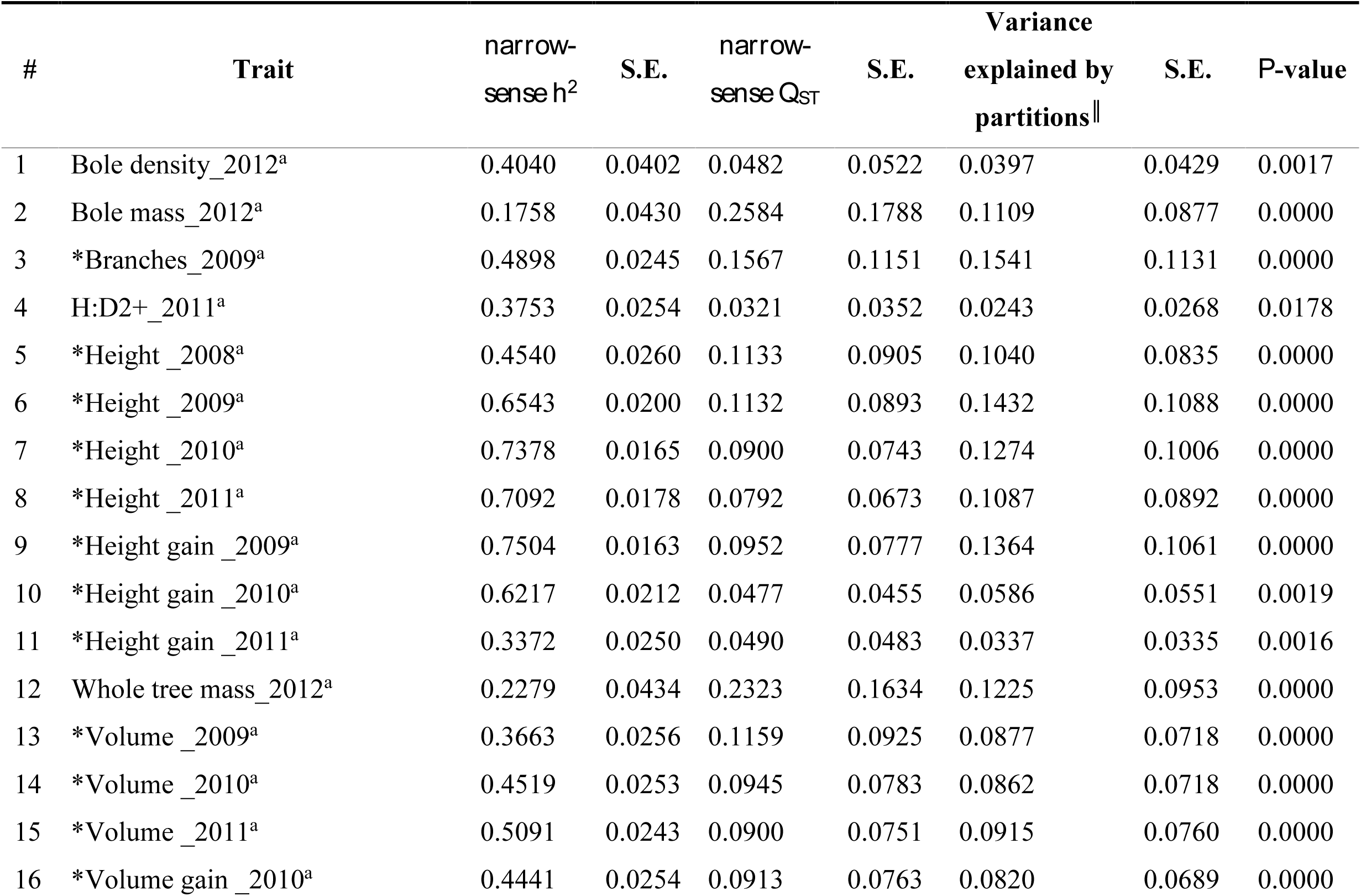
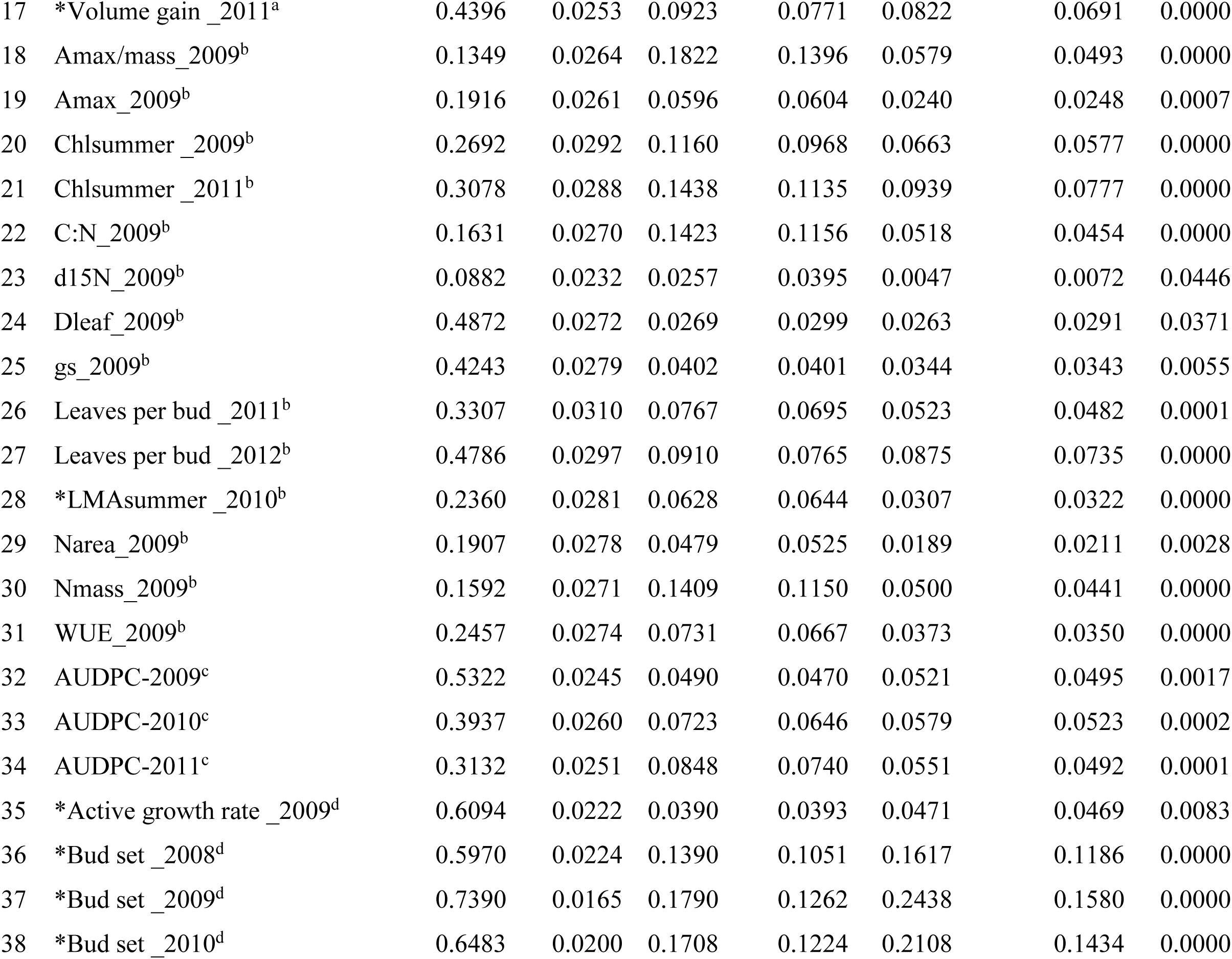
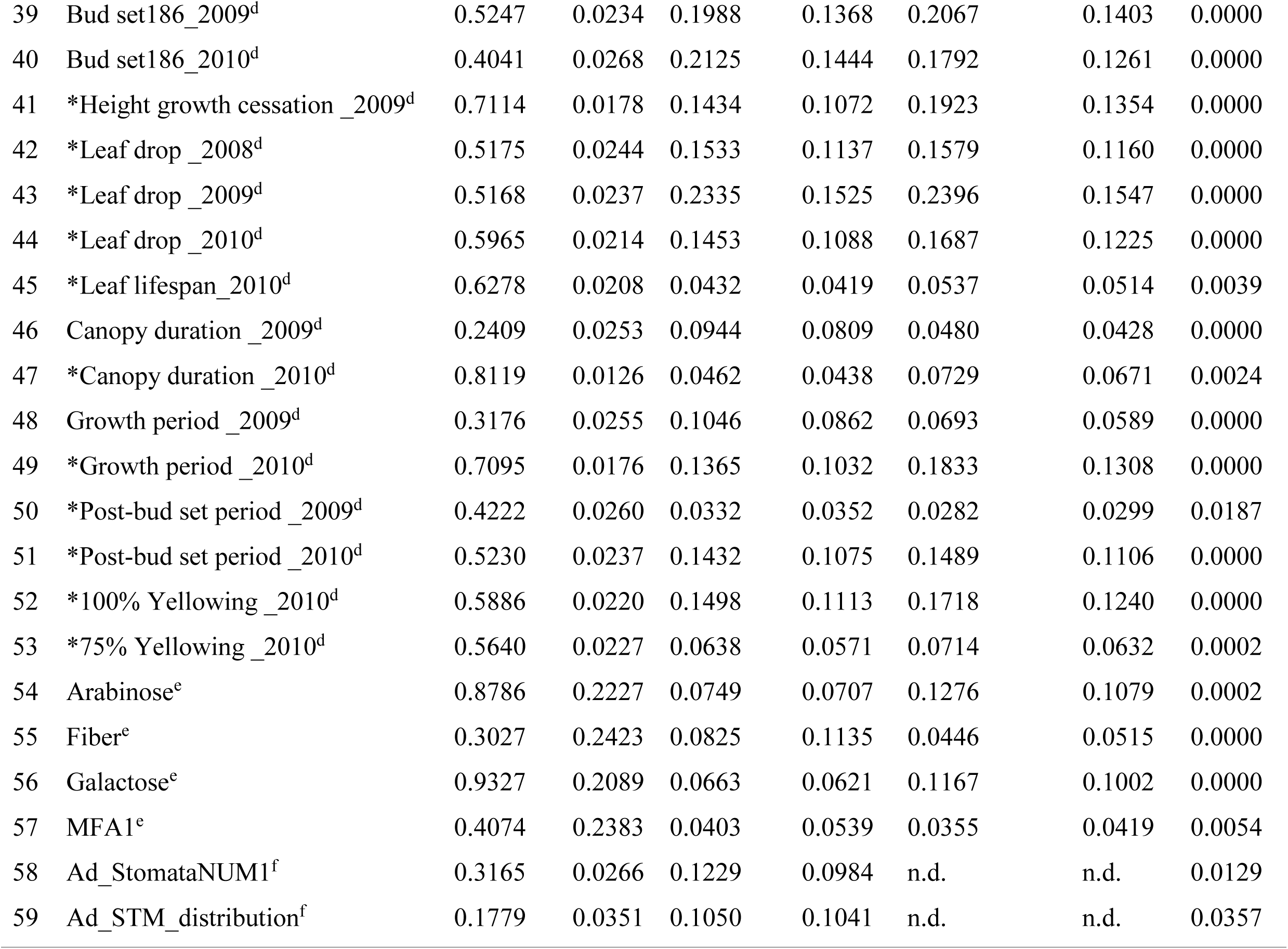
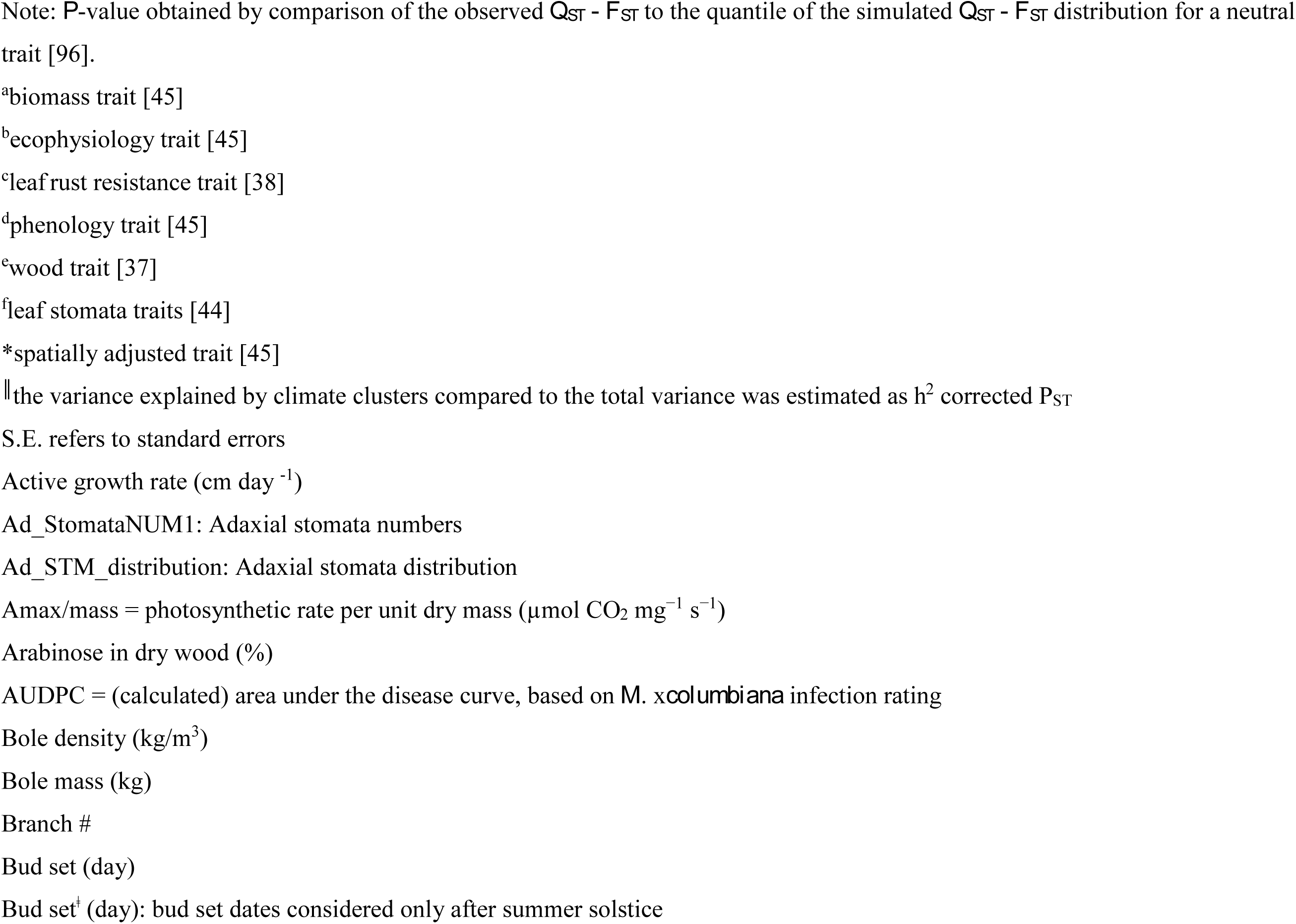
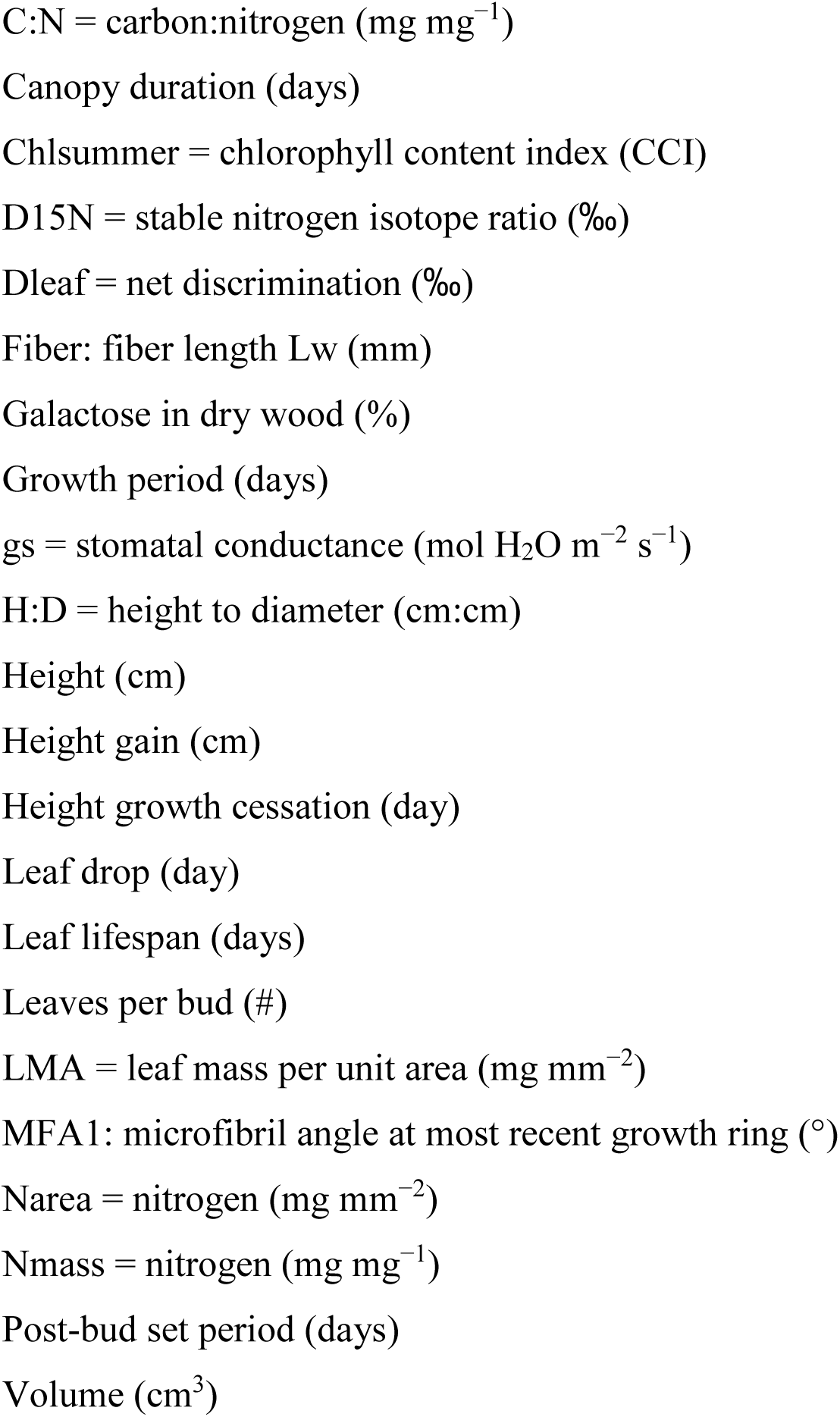
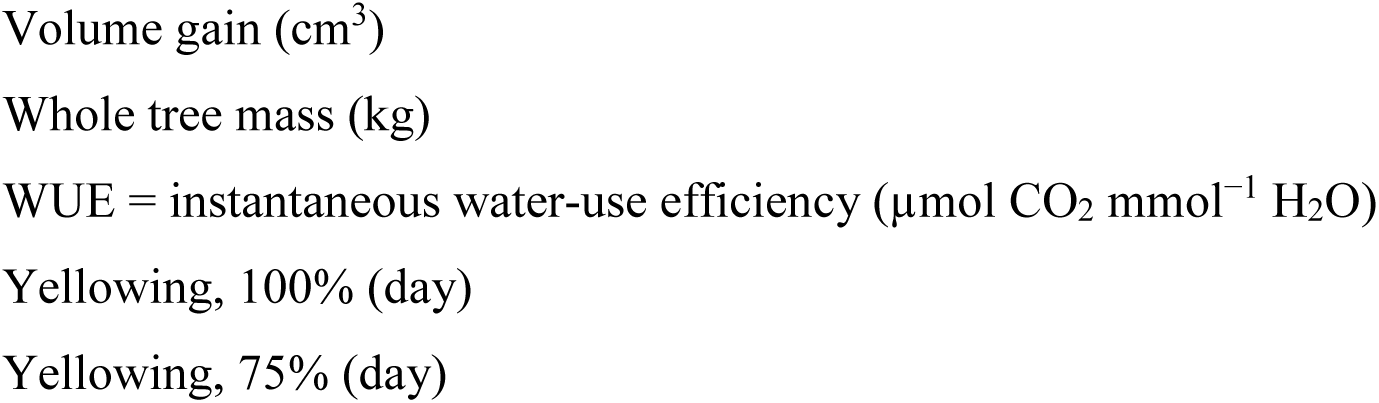
*h*^2^, *Q*_*ST*_, and *h*^2^ corrected *P*_*ST*_ of adaptive traits (*P*<0.05) Summary of 39 distinct *adaptive* traits of *P. trichocarpa* that diverged among different climate clusters (displayed are 59 tests for adaptation including tests for traits replicated in time, comprehensive results shown in Table S1)

### Identification of SNPs under selection

Using both unsupervised and climate-based SPA, a total of 1,468 SNPs were identified being under selection at a 5% cutoff for each method (Table S2). We also performed *F*_*ST*_ outlier analysis on climate clusters. While the mean *F*_*ST*_ value for the complete dataset (29,354 SNPs) was 0.0108, we obtained a mean neutral *F*_*ST*_ value (0.0078) after removing loci identified to be potentially under selection [41]. In the final analysis, all loci were tested against this neutral mean to identify a set of potential *F*_*ST*_ outliers relating to climate. Using 200k simulations in Fdist2, we identified 121 SNPs outside the 99% limits of the neutral distribution (Fig. S1) as potential candidates subjected to diversifying (positive) selection related to the four climate clusters. Among these, 88% of these climate-related ‘outliers’ were confirmed by allelic frequency correlation analysis with averages for climate variables within subpopulation (using multiple univariate logit regression models in SAM (α=0.05, Table S2)), 77 of these loci persisted across different selection scan scenarios employed (unsupervised SPA, climate-based SPA, and *F*_*ST*_ analysis based on population subdivision [36]), and 48 SNPs were retrieved using association genetics (see below) (Table S2). A comparison between Fdist and SPA testing gene dispersal and employing Moran’s test for spatial autocorrelation (Fig. 3) indicates, in general, the higher effectiveness of SPA to identify genetic selection signals under patterns of IBD.

**Fig. 3.**
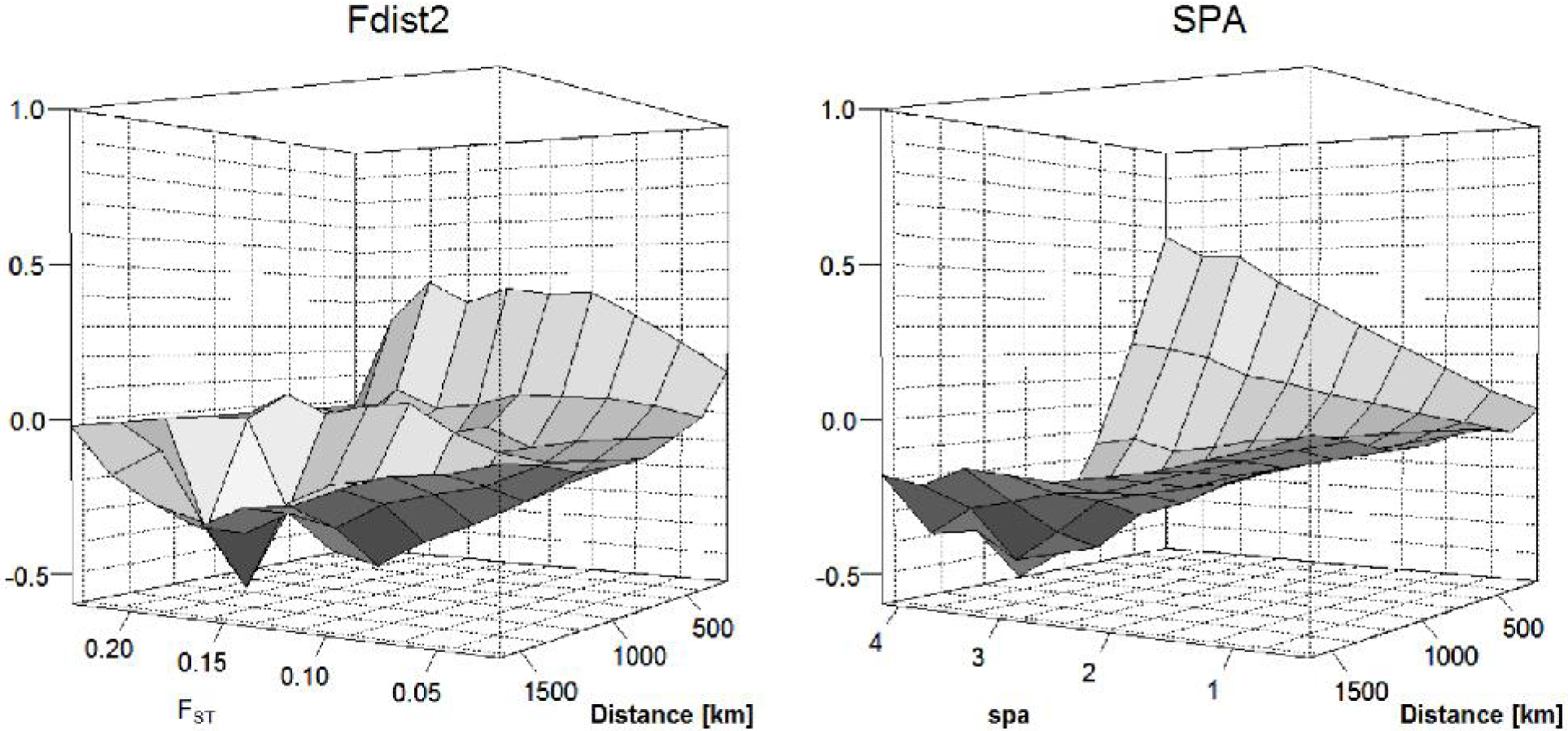
Comparison of two outlier detection methods (*F*_*ST*_, SPA) for their efficiency to identify genetic selection signals under isolation-by-distance (IBD). Gene dispersal was tested employing Moran’s test for spatial autocorrelation using 200km lags. (TIFF)

A significant accumulation of *F*_*ST*_ outliers was identified on chromosome 15 (Fig. S1). The extent of linkage disequilibrium (LD) between all 121 outlier loci is presented in Fig. S2. In general, we found that LD was not substantial between SNPs from different genes. Incomplete LD can be caused by the possibility that SNPs are close to but not in complete LD with the causal variants (here probably due to ‘tag SNP’ design of the SNP chip array [35]) explaining why the observed LD between diverged loci is generally low [42] One notable exception is two neighboring poplar genes (Potri.009G008600 and Potri.009G008500) initially annotated based on sequence homology to *Arabidopsis* genes as nitrate transporter types *ATNRT2:1* and *ATNRT2:4*, respectively. The allele frequencies of three SNPs and one SNP, respectively, in poplar orthologs of *ATNRT2:1* and *ATNRT2:4*, respectively, are strongly correlated to temperature (R^2^>0.9; *P*=0.05), while the remaining SNPs in both genes did not follow such a strong pattern (Fig. S2).

### SNPs under diversifying selection and associated with quantitative traits

To corroborate findings of candidate loci putatively under diversifying selection based on climate, we compared these results with SNPs uncovered by associations with adaptive traits (showing non-neutral *Q*_*ST*_). Among four GWAS studies in *P. trichocarpa*, a total of 619 SNPs had been identified with significant trait associations (at α=0.05): 410 with biomass, ecophysiology and phenology [39], 141 with wood property traits [43], 40 with *Melampsora* x*columbiana* resistance [38], and 28 SNPs related to leaf stomata variation [44].

We compared four different outlier analyses to identify selection signals in 29,354 SNPs. Most trait-associated SNPs for which we detected selection signals were associated with adaptive traits (89%, Table S2). The highest percentage of trait-associated SNPs in outlier analyses was found for climate-based *F*_*ST*_ outlier analysis (40% of the total number of outliers identified by the method; 48 SNPs), followed by geography-based *F*_*ST*_ outlier analysis (8%; 75 SNPs that were reported in [36], unsupervised SPA (5%; 75 SNPs), and SPA with climate as a covariate (3%; 37 SNPs). In total, selection signals were detected for 151 trait-associated SNPs with 44% overlap among evaluation methods. Interestingly, there was a lack of genome-wide correlation between selection and association signal (Fig. 4) and thus only dispersed association signals were detected among SPA selection signals (Fig. 5, Table S2). This result is probably a consequence of the structure correction methods employed in GWAS.

**Fig. 4.**
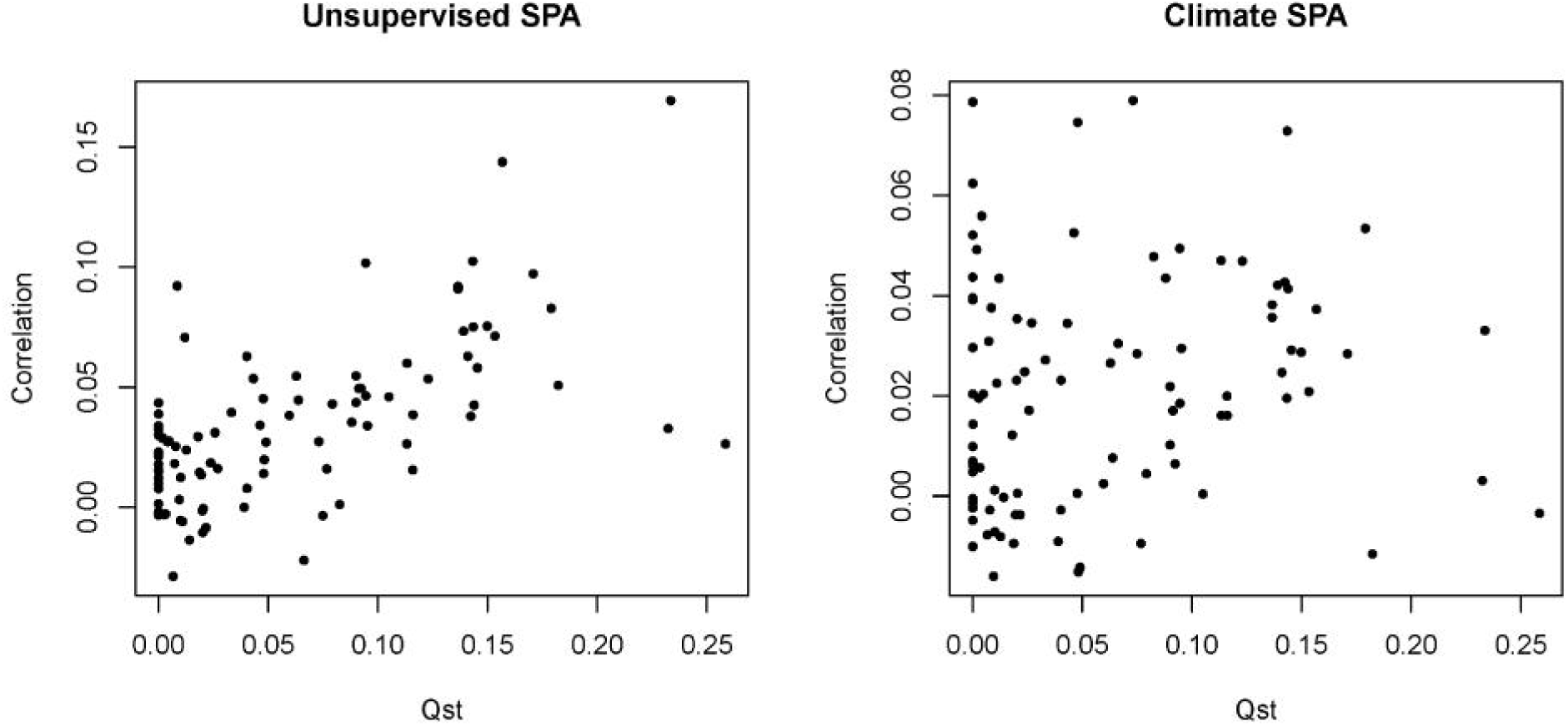
Genome-wide correlations between selection outliers and association signals based on 29k SNPs. Correlation of -log (*P*) versus spa was plotted against the trait’s *Q*_*ST*_. (TIFF)

**Fig. 5.**
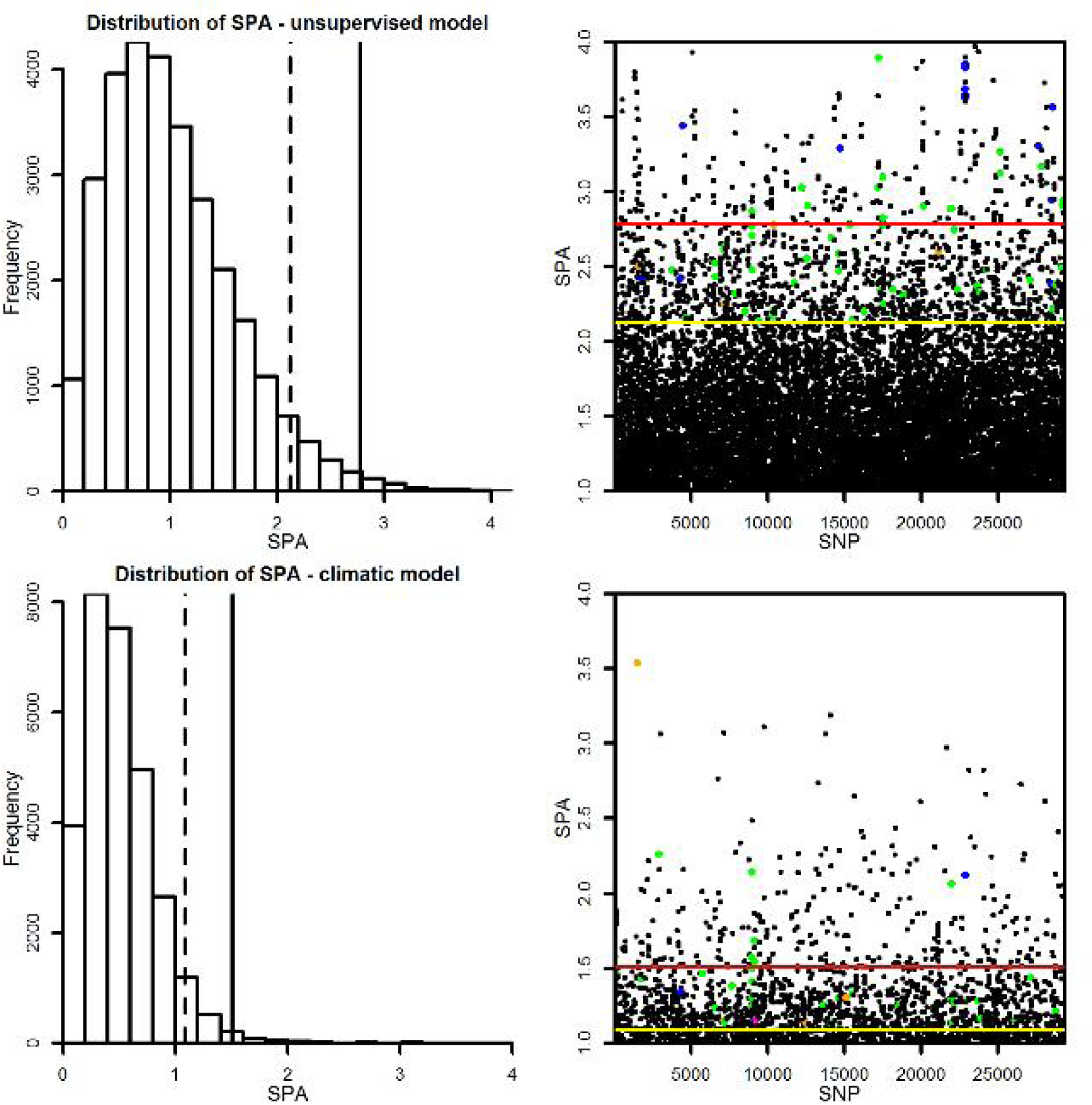
Individual SNPs under diversifying selection within genes mapping to quantitative trait variation. 5% cutoff: dashed and yellow lines; 1% cutoff: solid and red lines; ecology (biomass, ecophysiology, phenology, stomata) - green dots; wood properties (orange); rust resistance (blue) (TIFF)

We retrieved a number of unique but also shared SNPs among the different analyses (Fig.6). Shared SNPs were highest for climate *F*_*ST*_ (75%) and geography-based *F*_*ST*_ (72%). Unsupervised SPA had the highest number of unique SNPs among the four methods (51%). We found 118 SNPs associated with adaptive traits (significant *Q*_*ST*_) including 59 SNPs under diversifying selection shared among at least two outlier detection methods and 59 unique SNPs detected by climate *F*_*ST*_, climate SPA and unsupervised SPA, respectively (Table S3). A large number of SNPs (40%) that we identified as *F*_*ST*_ outliers using climate clustering were candidate SNPs from association studies (Table S2). The high number of trait-associated SNPs reflects both the polygenic nature of phenotypic traits (*e.g.*, c.200 for bud set, [39]) and linkage disequilibrium (LD) to a lesser extent. The highest number of climate-based *F*_*ST*_ outliers associated with adaptive traits was found on chromosome 15 (12 SNPs), identifying a genomic region where SNPs putatively under selection to local climate generally may be clustered (Fig. S1).

**Fig. 6.**
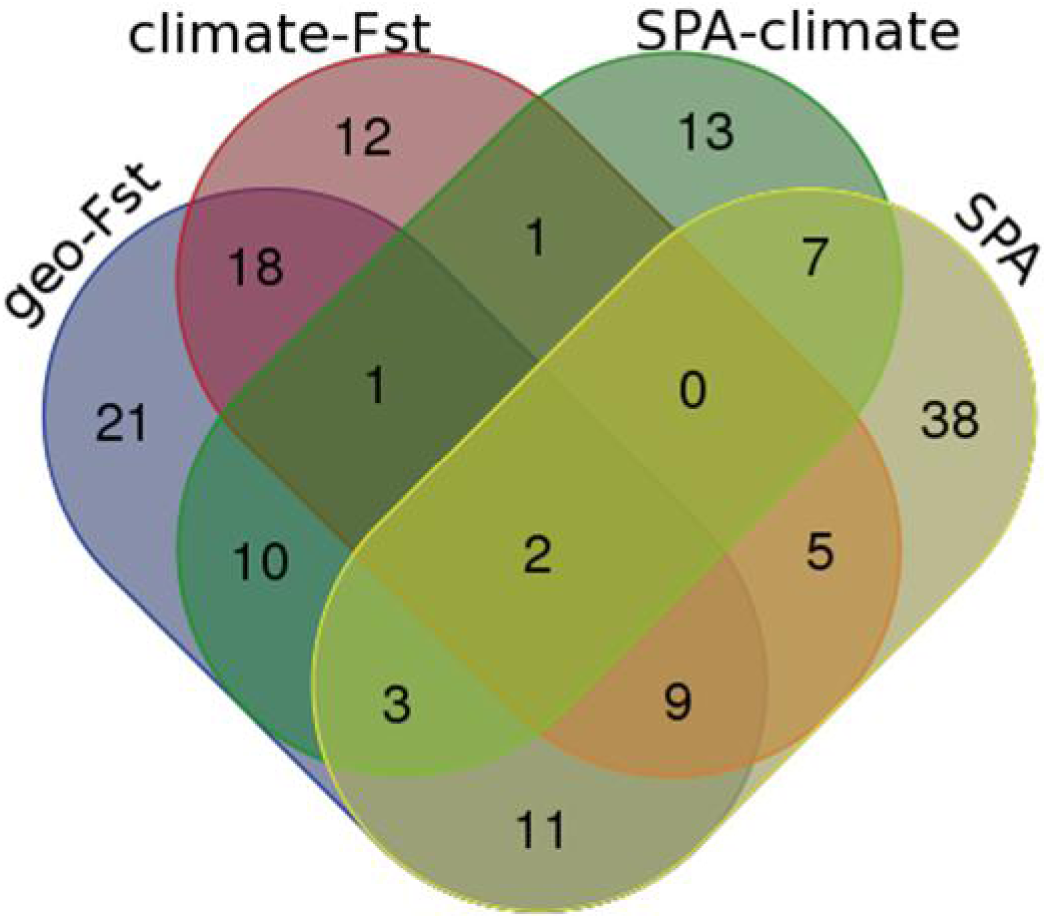
Venn diagram showing the numbers of unique and shared SNPs (totaling 151 trait-associated SNPs) among four different outlier detection approaches. *F*_*ST*_ using climate clusters, *F*_*ST*_ using geographical grouping, SPA analyses - with climate-based PCs incorporated as covariates and unsupervised, respectively. A subset of this information (118 SNPs) related to genetic polymorphisms associated solely with *adaptive* trait variation is provided in Table S3. (TIFF)

We found that SNPs under potential climate selection matching putative causal variants from association studies consistently mapped to non-neutral *Q*_*ST*_, *adaptive* traits (Table S1, Table S2). Only one SNP associated with wood traits (within Potri.009G006500 annotated as *FRA8* associated with fiber length, [43]) was among the *F*_*ST*_ outlier loci. Comparatively, phenology traits were the most complex *adaptive* traits from the high match between the total number of associated SNPs and the proportion of SNPs with allele frequencies significantly diverged among climate clusters (Table S2). In total, 118 SNPs were outliers under diversifying selection, associated with *adaptive* traits (significant *Q*_*ST*_), and with many SNPs putatively pleiotropic for functionally uncorrelated adaptive traits, such as autumn phenology, height, and disease resistance (Table S3). The 78 annotated poplar genes were largely derived from major gene functional group such as (1) transcription factors of several categories and (2) carbohydrate-related genes, but also transporters. Among these transporters, two poplar genes (Potri.009G008600 and Potri.009G008500) annotated based on sequence homology to *Arabidopsis* genes as nitrate transporter types *ATNRT2:1* and *ATNRT2:4*, respectively, were highly pleiotropic for several adaptive traits (Table S3).

## Discussion

### Evolutionary quantitative genomics

The main focus of our work involved identifying adaptive traits and their genetic basis in forest trees by employing both a quantitative genetics approach (*Q*_*ST*_ analysis) and population genomics [16] to uncover SNPs under strong selection (among c.29k tested genetic polymorphisms). Our analyses revealed that 53% of these traits produced significant *narrow-sense Q*_*ST*_ (Table S1) underscoring that such quantitative traits are very likely related to adaption to local climatic conditions [45].

This study uses SNP marker-inferred relatedness estimation (*i*.*e*. the ‘animal model’) to obtain *narrow-sense* estimates of heritability and *Q*_*ST*_ in wild populations [24]. The quality of genetic estimates using the ‘animal model’ approach largely depends on the accuracy of relationship coefficient estimates and are affected by: 1) number and quality of markers [46], 2) variance in actual relatedness [47], and 3) how well the relationship estimates reflect the segregation of causal variants [48] Our present study is based on extensive, genome-wide SNPs [35] which can provide high accuracy for both the relationship coefficients and the estimated genetic parameters. However, samples from natural tree populations are subject to intensive gene flow (outcrossing) and generally show low levels of relatedness which can negatively affect heritability and *Q*_*ST*_ analyses.

Heritability is usually dependent on the population sampled (*i.e*. the observed allele frequency differences) and thus, can differ for smaller sampling sizes and/or specific sampling areas (*e.g*., central vs. marginal regions of species distribution). Heritability estimates taken across a greater coverage of the species distribution are more likely to reflect evolutionary history of the traits (stabilizing vs. diversifying selection) rather than the effects of population subsampling. Sufficient variance in the actual relatedness is also required to reveal heritability in wild populations [47], although heritability, and indirectly, *Q*_*ST*_ estimates, can suffer from the inability to separate the pure additive genetics from environmental effects, specifically when relatedness is lacking. Thus, the presence of LD between markers and causal variants (QTLs) is crucial to recover the genetic parameters with sufficient precision. In the case of traits under diversifying selection, the additive genetic variance estimates (such as *narrow-sense* heritability) may also include a substantial QTL covariance component, in addition to the pure genic variance. This is especially the case when many QTLs follow the same cline, and can further extend the additive genetic variance when the QTLs interact (*i*.*e*., epistasis) [49] unless the epistasis is accounted for in the model [50]. Thus, heritability estimates for traits under diversifying selection (Table 1) may be upwardly biased (see below).

Heritability estimates are often interpreted as the capacity for adaptive evolution. In addition, epistatic interactions, specifically, the directional epistasis, have major effects through altering the genetic background (both, the additive genetic variances and the covariances, *i*.*e*. the allelic frequencies but also their effects) [51]. Hemani *et al*. (2013) outlined that for traits under selection, high levels of genetic variation are maintained and the traits evolve more slowly than expected, yet this could be attributed to high epistasis in traits under strong diversifying selection [42].

### Selectively non-neutral genetic variants underlying traits adaptive to climate

Overall, the number of *F*_*ST*_ outlier SNPs underlying an adaptive trait correlated well with the total number of candidate SNPs associated with that trait (r=0.625, P=0.0005). Yet, the majority of trait associated SNPs were not *F*_*ST*_ outliers (Table S2) and appeared to be unresponsive to selection for different climatic conditions, especially for phenology traits such as bud set, leaf drop or growth period. A previous simulation study suggested that differentiation in candidate loci is limited for complex traits in forest trees (*i*.*e*., their *F*_*ST*_ values are similar to neutral values), despite their strong adaptive divergence among local populations (high *Q*_*ST*_), due to large population sizes and high levels of gene flow [52]. Thus, highly polygenic adaptation (as observed in complex genetic traits) will not show sufficient allele frequency differentiation such that climatic clines in SNPs of candidate genes can be exhaustively detected.

We modelled the spatial structure of genetic variation using SPA (addressing gene flow under IBD), and SNPs identified via SPA were compared against GWAS-identified SNPs, climate-related *F*_*ST*_ outliers and geography-informed *F*_*ST*_ outliers. The majority of SNPs with steep allele frequency clines (based on unsupervised SPA) uncovered allele frequency correlations with the north-south cline (Table S2). We noted that enrichment for particular genes, such as circadian rhythm/clock genes, was found in PC1 (a north-south population structure) [45] and that SNPs of these genes were among the highest ranked in SPA. Nonetheless, associations of circadian rhythm clock genes with strong correlations to environment were largely missing among the identified genetic associations for phenology traits (discussed in McKown *et al*. [39]). The interplay of IBD and natural selection was lost by the necessary structure correction in GWAS, however, evidence from gene expression or gene regulation that is also highly correlated with the trait under question might be possible to retrieve such SNPs of putative importance (Anonymous, [53]).

The presence of IBD in *P. trichocarpa* underscores the larger issue for investigating wild populations with quantitative genetics and population genomics approaches as IBD can confound population structure, association mapping, and outlier analyses. The power to detect local selection depends on several factors, including selection strength, the presence of distinct types of microenvironment heterogeneity, and the distance of gene dispersal compared to the overall spatial scale [54]. In our case, as the observed gene dispersal is ∼500 km (Fig. 3) and sampling is also discontinuous (Fig. 1), this does not allow us to perform *F*_*ST*_ analysis on arbitrarily defined local populations because it will be more difficult to separate the stochastic noise (drift, migration) from the selection signal in smaller scale population subsampling leading to an excess of false positives [54]. Yet, selection pressures can differ along environmental clines. Thus, *F*_*ST*_ outliers should be investigated on the largest scale possible following the spatial distribution of the environment in order to identify spatial genetic structure. Nevertheless, IBD in wild populations will create some compromised statistical power in detecting local adaptation using specific pairs of populations that is unavoidable (Fig. 3).

### Polygenic and pleiotropic adaptation relating to climate

Our climate clustering partitioned the study population into four large, evenly-sized groups of individuals lending robustness to SNP detection even for lower frequency (recent) variants. In our study, the top two SNPs among climate related *F*_*ST*_ outliers showed strongest associations to climate partitions according to SAM analysis [Potri.010G250600 (*MSR2*/ *MANNAN SYNTHESIS RELATED 2* implicated in carbohydrate metabolism) and Potri.010G254400 (transporter *ATGCN4*) (Table S2)]. In addition, six genes that harboured climate-related *F*_*ST*_ outlier SNPs have been identified as candidates for bud set in previous studies ([55]; [56]), yet these loci were not associated with bud set in our GWAS study ([39]; Table S2), possibly through implementing the conservative population structure correction term in GWAS. Nevertheless, these genes may represent additional candidates for bud set, including Potri.003G218900 (*ACD1-LIKE*), Potri.009G015100 (senescence-associated family protein), Potri.014G170400 (*XERICO*), Potri.015G012500 (*IQ-domain 21*), Potri.018G015100 (chloroplast nucleoid DNA-binding protein), and Potri.019G078400 (leucine-rich repeat transmembrane protein kinase) (Table S2).

Evidence is emerging that for perennial trees to effectively sense short day signals, *i*.*e*. critical day length in autumn phenology [57], a temperature optimum is required and genetically pre-determined by the local climate of the individual’s origin [58]. Allele frequencies for most of the SNPs that both associated with bud set and diverged among the climate clusters showed strong regression on the mean temperature variation of the climatic clusters (R^2^ up to 0.94; Table S2). A critical role for temperature, rather than precipitation, on bud set has also been found in *Picea* [12]. For autumn phenology, elevated temperatures can either accelerate or delay growth cessation depending on species or ecotype ([59]; [60]), but under climate warming, the overall effects on phenological timing in forest trees is unknown.

SNP allelic frequencies within both nitrate transporter genes *ATNRT2:4* and *ATNRT2*:1 were strongly aligned with temperature variation (R^2^∼90%) in *P. trichocarpa*. Moreover, these SNPs were pleiotropic for multiple autumn phenology traits, height, and leaf rust resistance (Table S3). Nitrate transporters are generally important in plants, as nitrate is the main nitrogen source required for synthesis of nucleic and amino acids. Therefore, a regulation of nitrate distribution is crucial to modulate growth (biomass acquisition) in response to temperature or light conditions ([61]; [62]). Interestingly, there are only two poplar representatives within a phylogenetic sub-clade of NRT2 that is populated by as many as five Arabidopsis sequences (*ATNRT2.1/2.2/2.3/2.4/2.6*). This implies that a deletion event occurred in this clade whose functional significance remains elusive to date [62]. Phylogenetic reconstruction coupled with gene expression analysis point at neo/subfunctionalisation of the two poplar nitrate transporters for long distance nitrate transport from roots, wood to leaves [62]. This acquisition of novel expression pattern and loss of the ancestral expression pattern demonstrates the signature of adaptive evolution in functional diversification in paralogous gene pairs [63].

In addition, our results revealed that adaptive genetic variants within both poplar nitrate transporters were also associated with leaf rust resistance ([38]; Table S3). In *Arabidopsis*, loss of function of *ATNRT2.1* primes salicylic acid signaling and *PR1* up-regulation [64]. In poplar leaf rust inoculations, both *PTNRT2.4* and *PTNRT2.1* are strongly down-regulated in incompatible interactions, while no expression change is apparent in compatible interactions (J. La Mantia, personal observation). The identified nitrogen transporters might be important in nitrogen storage and nitrogen remobilization to recycle nutrients during the progression of leaf senescence [65]. They may also function in down-regulation of nitrogen assimilation during seasonal remodeling of tree phenology related to growth cessation induced by short photoperiods ([66]; [67]) and/or temperature [58]. The effect of temperature on rust aggressiveness is noted [68] and the climatic conditions which form a conducive environment for rust infection and disease duration likely provide a strong adaptive selection toward resistance.

Pectin esterase gene Potri.012G014500 (SNP scaffold_12_1811250) represents another example for which significant associations with climate (here: temperature) and several adaptive traits were found (Table S2, Table S3). In fact, the allelic effects of this SNP related to hypostomaty also related to less rust infection ([45]). This is an illustrative example regarding the tradeoff between carbon gain and pest resistance under favourable climatic conditions relating to pathogen pressure ([45]).

## Conclusions

The high adaptive potential of tree populations is considered the result of positive effects of long-distance gene flow based on its interactions with divergent selection across the contrasting environments [69], while local adaptation in forest trees with regards to climate-related traits is polygenic and recent [70]. For instance, interactions between temperature and photoperiodic cues were shown to influence bud set for short-term acclimation in poplar [58]. By combining quantitative genetics and population genomics analyses, our study contributes to an enhanced understanding of the molecular basis of adaptation to different local climate in an undomesticated perennial species (*P. trichocarpa*). The key findings provided SNPs whose allelic frequencies were most diverged among populations from different climate clusters and these SNPs tended to be associated with mapped genes underlying phenotypic variation. This phenotypic variation itself diverged among the different climate clusters. Our study dissected the influence of climate (specifically, temperature and precipitation), yet much of the variation in phenology is also attributed to photoperiod ([71]; [72]; [45]). The tight photoperiodic control of traits such as bud set, height growth cessation, and leaf senescence ([73]; [74]; [59]) is crucial both for resistance to cold temperatures and maximization of the growing season, particularly in trees originating from high-latitude and/or high elevation provenances ([75]; [56]). While we tested the influence of climate on the variation of other traits in *P. trichocarpa*, such as wood and biomass, we consider other local factors, such as soil condition (pH and minerals), soil/root microbial diversity, groundwater, and other ecological interactions also of potential importance. Reciprocal transplants will be necessary to elucidate the effects of gene × environment plasticity on the expression of traits with spatially heterogeneous selection [76], but can focus on specific genes identified through a combined quantitative genomics analysis, such as the one proposed here. Forthcoming research can also scale trait-to-performance mapping in known pedigrees for the assessment of SNP effects on fitness [77]. These findings will have important implications for the future management of natural forests, acting to guide efforts in facilitated adaptation to climate change via measure such as assisted gene flow [78].

## Materials and Methods

### Collection, genotyping, and phenotyping of *P. trichocarpa*

Plant material was collected from a population of 433 *P. trichocarpa* Torr. & A. Gray genotypes growing in a common garden. These genotypes came from 140 unique geographic locations spanning two thirds of the species’ range (44-60°N, 121-138°W) ([79], Fig. 1). Originally collected by the BC Ministry of Forests, Lands and Natural Resource Operations, individual genotypes were grown in two common gardens, Surrey, BC and Totem Field, University of British Columbia, BC. Genotypes were replicated across the two field gardens and the Totem Field individuals (established in 2008 [80]) were clonal propagations from Surrey site individuals (established in 2000 [79]).

Trees were genotyped using an Illumina iSelect array with 34,131 SNPs from 3,543 candidate genes designed for *P. trichocarpa* [35]. The characteristics of the poplar genome and array development are outlined in [35]). Briefly, the SNP array was designed to include genes of known importance (*i*.*e*. candidate genes) or genes based on expression analyses. Because of the rate of linkage disequilibrium (LD) decay in *P. trichocarpa*, between 67 – 134k SNPs would be required to include all common variants throughout the genome at LD=0.2 (assuming a 403 Mb assembled genome length and an average of 3–6 kb for r^2^ between common variants to drop to 0.2). Therefore, some SNPs were selected as representative SNPs to “tag” genes and genetic regions with high LD, and thus represent a group of SNPs (the haplotype). For this study, we further filtered array SNPs for: i) minor allele frequency (MAF) <0.05, ii) >10% missing data, and iii) Illumina’s GenTrain score <0.5, thereby reducing SNP numbers to 29,354. This filtering is not biased towards higher frequency SNPs (*i*.*e*. older variants established at much higher frequencies within the population over time) as a wide distribution of allele frequencies (MAF>0.05) was considered for the analysis.

Phenotyping of genotype accessions within the common gardens and climate of origin data were obtained from previously published work (for full phenotyping details, see [38]; [37], [45]). In brief, phenology, ecophysiology, biomass [45], leaf stomatal anatomy [44] and leaf rust (*Melampsora* x*columbiana*) resistance traits [38] were repeatedly measured from accessions planted at the University of British Columbia’s research field through replication in space (clonal ramets) and in time (measurements across years). Wood chemistry and ultrastructure traits were measured from wood cores of the nine-year-old ortets representing the same genotypes and growing in Surrey [37].

### Assessment of population structure

Since forest tree species usually have extensive geographic ranges, exhibit extensive gene flow and have low levels of population stratification [81], we investigated whether the genetic variability due to non-random mating in our population was caused solely by isolation-by-distance (IBD), reflecting the large geographical distribution of our sample (cf. [36]), or also by natural barriers causing local genetic clusters. We performed spatial principal component analysis (sPCA) by using the “spca” function implemented in the R package “adegenet” [82] which is a spatially explicit multivariate analysis accounting for spatial autocorrelation processes and patterns of genetic variation. A K-nearest neighbours method with K = 10 was used as connection network. Positional information for each genotype were transformed into Universal Transverse Mercator (UTM) coordinates using “convUL” in the R package “PBSmapping” [83]. Due to the occurrence of multiple genotypes with identical geographical coordinates (*i*.*e*. trees collected at the same latitude/longitude), we randomly selected a single genotype representing a geographical region (out of the total 140 locations). Eigenvalues for principal components from sPCA provided a cumulative picture about contributing factors, including the genetic variance and the spatial autocorrelation (through Moran’s I, see below). Large positive eigenvalues reflect the importance of the proportion of the genetic variance along with a strong positive autocorrelation in the global pattern (*i*.*e*. IBD), while large negative eigenvalues indicate the importance of the proportion of the genetic variance along with negative autocorrelation indicating the existence of discrete local genetic clusters.

We used the "global.test" and "local.test" functions in the "adegenet" package to infer the statistical significance of each type of genetic structure. These functions are based on a spectral decomposition of the connection matrix into Moran′s eigenvector map and test for association of those eigenvectors from Moran′s eigenvector map with Moran′s I [82]. To investigate gene dispersal, we employed a Moran I test for spatial autocorrelation ([84]; [54]). Moran’s I coefficients were investigated in 200 km spatial lags and the analysis was performed using “moran.test” in the “spdep” R package [85]. Moran’s I coefficients were estimated as follows:

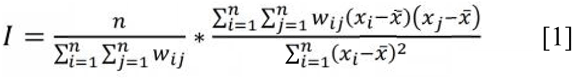

where *n* is the number of populations (*i*.*e*. unique geographical locations), *W*_*ij*_ is weight set at 0 or 1 depending on whether populations are considered neighbours in each 200 km lag test, *x*_*i*_ is the allele frequency in the i^th^ population, and 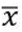 is the allele frequency across all populations.

### Climatic zone clustering of *P. trichocarpa*

Since our initial investigation of population structure with sPCA indicated the presence of only one global structure consisting of IBD and lack of local discrete clusters, any marker-based inference about genetic clusters might be highly unreliable [86]. Therefore, we established population differentiation on the basis of climate envelopes ([12]). Clusters of individual genotypes were defined using climate of origin measures (*i*.*e*. independently of the genetic data). Climate variables were obtained using ClimateWNA [87] and included mean annual temperature (MAT; °C), number of frost-free days (NFFD), and mean annual precipitation (MAP; mm). Climate data were based on positional information (latitude, longitude, elevation) and 1971-2002 Canadian Climate Normals [45]. Using K-medoids clustering and the Calinski-Harabasz criterion [88], we split the study population into four groups with relatively balanced sample sizes of 87, 103, 142, and 101 representing climate classes #1-4, respectively. Clusters generally followed the western North American coastline inwards (Fig. 1*a* & *b*).

### Genetic differentiation in quantitative characters among populations defined by climate clustering

We tested phenotypic characteristics in *P. trichocarpa* for their adaptive potential (Table S1). For *Q*_*ST*_ – *F*_*ST*_ comparisons, *Q*_*ST*_ values among the identified climate-related population groups were first estimated for each trait following [89] and [24], respectively.

The *narrow-sense Q*_*ST*_ was estimated by computing the variance components using the ‘animal model approach’ [90] following:

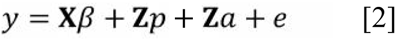

where *β* is a vector of fixed effects (intercept), *p* and *a* are vectors of random climate cluster and individual tree additive genetic effects, **X** and **Z** are incidence matrices assigning fixed and random effects to measurements in vector y, the cluster effects are following 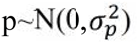 where 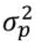 is the cluster variance, individual tree additive effects are following 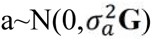 where 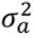 is the additive genetic variance and **G** is the realized relationship matrix [91], using 29,354 SNPs estimated in R package “synbreed” [92] as follows:

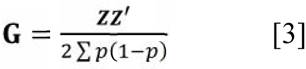

where **Z** is **M**-P, with M the marker matrix with genotypes recoded into 0, 1 and 2 for the reference homozygote allele, the heterozygote and the alternative homozygote allele, respectively, and with P the vector of doubled allele frequency; e is the vector of random residual effects following 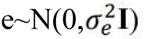 where 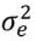 is the residual variance and **I** is the identity matrix. The *narrow sense Q*_*ST*_ was estimated as follows:

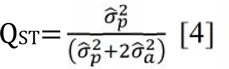

where 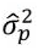 and 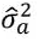 are the estimates of cluster and additive genetic variance representing among- and within-group trait variances attributable to additive effects.

The measurements of all ecology and disease traits using clonal ramets (*i*.*e*. replication) enable estimating *broad-sense Q*_*ST*_ directly without the use of any relationship matrix, while *narrow-sense Q*_*ST*_ estimation was based on variance components estimated in the mixed linear model considering the realized relationship matrix [91] as in equation 2. The model is identical to equation 2 where the variance components for *broad-sense Q*_*ST*_ were estimated in the model considering a as the vector of clonal genotypic values following 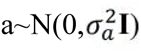 where 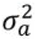 is the total genetic variance (including both additive and non-additive component) and *e* as the vector of ramet within clone effects following 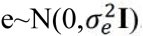. Then, the computed *Q*_*ST*_ values for each trait were compared to the average population differentiation estimate (*F*_*ST*_) strictly based on neutral markers (see below) allowing inferences about trait evolution based on selection or genetic drift (neutral trait), [93].

*Narrow-sense* heritability (*h*^2^) was based on variance components estimated in the mixed model as follows:

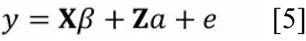

where *β* is the vector of fixed effects (intercept and cluster) and a is the random vector of additive genetic effects following the description of equation 2. The *narrow-sense* heritability was estimated as follows:

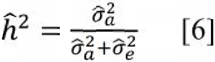

where 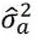 and 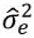 are estimates of additive genetic and residual variance, respectively. The phenotypic *Q*_*ST*_ (*i*.*e*. *P*_*ST*_) ([89]; [24]) was estimated as follows:

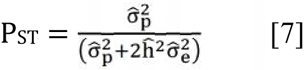

where 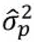 and 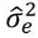 are estimates of cluster and residual variance representing among- and within-population variances, respectively, and 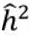 is the heritability estimated according to [37]. The variance components were estimated in ASReml software [94] using the mixed linear model following:

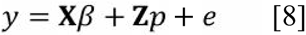

where *β* is the vector of fixed effects (intercept) and *p* is the vector of random cluster effects, the effect of individuals within cluster is found within the error variance.

### Identification of non-neutral SNPs and quantitative traits divergent among climate clusters

To identify SNPs putatively under selection and also associated with adaptive traits ([38]; [43]; [39]), we performed: 1) *F*_*ST*_ outlier analysis (using Fdist2) employing the same climate clusters as for *Q*_*ST*_ analysis, 2) unsupervised spatial ancestral analysis (SPA), and 3) SPA with climate as a covariate. Additionally, we compared our results with *F*_*ST*_ outlier analysis (using Fdist2 and BayeScan) that were reported in [36] using 25 topographic units separated by watershed barriers within the geographic area from Central Oregon, USA (44.3°N) to northern BC, Canada (59.6°N)).

*F*_*ST*_ values for SNPs were calculated among the four climate clusters (for definition and calculation, see above). We implemented the Fdist2 program within the LOSITAN project [41] for SNP *F*_*ST*_ outlier detection. Fdist2 compares the distribution of *F*_*ST*_ values of sampled loci to the modeled neutral expectation of *F*_*ST*_ distribution using coalescent simulations [9]. We employed the infinite alleles mutation model (as we investigated SNPs), a subsample size of 50, and ran 200k simulations. *F*_*ST*_ values conditioned on heterozygosity and outside the 99% confidence interval were considered candidate outliers.

Since *P. trichocarpa* populations have known structure related to IBD ([36] and this study), we applied spatial ancestral analysis (SPA), a logistic regression-based approach [86], to detect SNPs with sharp allelic frequency changes across geographical space (implying candidates under selection). The unsupervised learning approach (using only genomic data) was employed to obtain SPA statistics. In addition, we tested SPA including the first two principal components (PCs) based on climate variables (explaining 91% of the variance) as covariates to determine individuals’ location based on allele frequencies related to MAT, NFFD, and MAP climate components.

We investigated correlations between the outlier SNPs (based on climate clusters) and the environmental variables that defined the established climatic clusters (Fig. 1). Subpopulation averages for MAT, NFFD, and MAP were tested for correlations with SNP allele frequencies employing multiple univariate logistic regression models with the spatial analysis method (SAM; [95]). The significance of correlations was assessed using three independent statistical tests (likelihood ratio and two Wald tests) implemented in SAM and applying an initial 95% confidence interval for the statistical tests. We used the Bonferroni correction method (α=0.05) for multiple testing resulting in *p*<6.887052*10^-5^ for 726 tested models (242 alleles, three variables). Only those correlations that remained significant after Bonferroni correction for each of the three test statistics (*i*.*e*. the likelihood ratio and the two Wald tests) were retained.

Finally, we compared observed *Q*_*ST*_ values with the simulated distribution of *Q*_*ST*_-*F*_*ST*_ values for a neutral trait using previously provided R scripts [96]. In brief, a range of possible demographic scenarios was tested simulating the distribution of *Q*_*ST*_ values based on mean *F*_*ST*_ for neutral markers and mean *Q*_*ST*_ for neutral traits ([97]; [98]). For a neutral trait, the expected QST was estimated based on 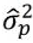 (*i*.*e*., measured within-population variance; see above) and 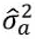 (*i*.*e*., expected between-population variance) given in equation 4. The distribution of 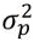 values was based on 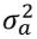 and the observed *F_ST_* values of 29,233 SNPs present (total number reduced by removing outliers) within the simulated *neutral* envelope of *F*_*ST*_ values (*F*_*ST*_ outlier analysis) with *Q*_*ST*_ replaced by the *F_ST_* in equation 4. *P*-values were obtained by testing whether the null hypothesis that the estimated *narrow-sense Q*_*ST*_ for each tested trait is statistically equal to the expected *Q*_*ST*_ for a neutral trait [96].

### Marker-trait association mapping

In previous analyses of marker-trait associations in *P. trichocarpa*, confounding effects of population stratification were adjusted using principal component analysis ([38]; [43]; [39] and a **Q** matrix population structure correction [39]. Phenological mismatch within the common garden can confound trait values [45], thus, association analyses included “area under the disease curve” resistance measures with adjustment for bud set [38] and all ecophysiological traits that were measured prior to bud set [39]. The Unified Mixed Model (a modification of the generalized linear model) was employed for marker-trait association mapping and is fully described ([38]; [43]; [39]). While necessary, the adjustment for confounding, cryptic genetic structure in the association analyses may have reduced the statistical power to detect associations. This is particularly problematic in species whose distribution is mainly along a one-dimensional cline or for which differentiation in ecological traits covaries with the species demographic history ([13]; [45]). Furthermore, the GWAS results may be biased towards common variants or variants with the greatest effects. This is related to the size of the SNP discovery panel (34k) [99] and the power to detect significant associations given the tested population sizes (334-448 individuals). As whole genome sequencing and phenotyping of thousands of genotypes would be required to comprehensively uncover the genetic architecture of complex traits, we consider the GWAS results informative but not exhaustive.

## Acknowledgements

The authors thank Dr. Julien Prunier for help with ‘Spatial analysis method’ software. This work was supported by Genome British Columbia Applied Genomics Innovation Program (Project 103BIO) and Genome Canada Large-Scale Applied Research Project (Project 168BIO), funds to RDG, RCH, JE, SDM, CJD, and YE-K.

## Author contributions

Conceived and designed the experiments: YE-K, RDG, RCH, JE, SDM, CJD, Performed the experiments: IP, ADM, JL, Analyzed the data: JK, IP, Contributed reagents/materials/analysis tools: PI, Wrote the paper: IP, JK, YE-K.

## Supporting table captions

**Table S1.** Comprehensive population differentiation estimates and *h*^2^ corrected *P*_*ST*_ for *P*. *trichocarpa: broad-sense* and *narrow-sense Q*_*ST*_ for 58 distinct field traits; *Q*_*ST*_*1* and *narrow-sense Q*_*ST*_ (*Q*_*ST*_*2*) estimates for 16 wood traits. (XLS)

**Table S2.** Comprehensive summary table of all SNP detection results from GWAS [ecology [39]; rust [38]; stomata [44]; wood [43]] and outlier analysis (geographic *F*_*ST*_ [36], this study: climate *F*_*ST*_, unsupervised SPA, climate SPA) for the black cottonwood population (presented in Fig. 1) and using the 34k SNP chip [35]; adaptive traits (significant *Q*_*ST*_) are in bold. In red and dark blue are 1% cutoffs (spa=2.78025 and spa=1.50795), in orange and light blue are 5% cutoffs (spa=2.12467 and spa=1.08868) in unsupervised SPA and climate SPA analyses, respectively. (XLSX)

**Table S3.** List of 118 SNPs associated with *adaptive* traits (significant *Q*_*ST*_ for at least one associated trait) including 59 SNPs under diversifying selection shared among at least two outlier detection methods and 59 unique SNPs detected by climate *F*_*ST*_, climate SPA and unsupervised SPA, respectively. Comprehensive results are provided in Table S2. (XLS)

### Figures

**Fig. S1.**
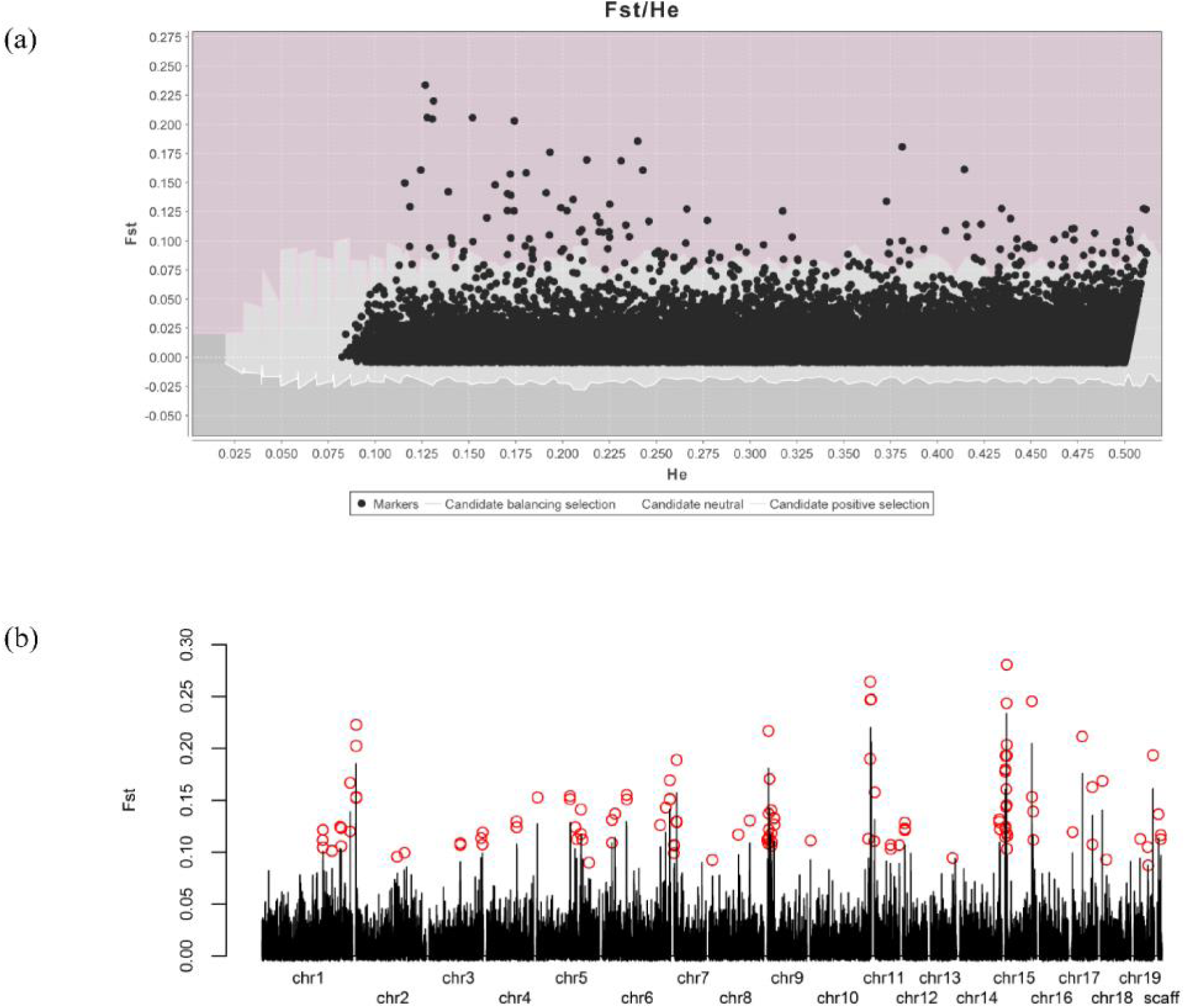
F_ST_ outlier loci detection in *P. trichocarpa* and distribution of outliers along the poplar chromosomes. Caption: (a) F_ST_ outlier loci detection and distribution of empirical *F*_*ST*_ estimates conditioned on expected heterozygosity (H_E_). The envelope of values corresponding to neutral expectations at 99% CI level (with mean F_ST_=0.0078), solid line, was constructed with the infinite allele model according to (Beaumont & Nichols, 1996) (b) Distribution of the empirical F_ST_ estimates along the 19 poplar chromosomes and additional scaffolds (abbrev: scaff); the 121 identified outlier loci are indicated by red circles above their F_ST_ value bars. A goodness-of-fit test assuming a uniform distribution was performed to test whether the observed frequencies of ‘outlier loci’ along the 19 poplar chromosomes differed significantly from the expected value. Following the rejection of the null hypothesis (chi-square = 81.98 df = 18, *p*-value = 3.85e-10), we declared ‘outlier loci hotspots’ if the number of outliers at a given chromosome was equal or above the maximum value (*i*.*e*., 20) for assessed outlier clusters from a randomly generated data set using the 118 outliers found across the 19 chromosomes, and running 1,000 replicates, which identified significant clustering of outliers on chromosome 15.

**Fig. S2.**
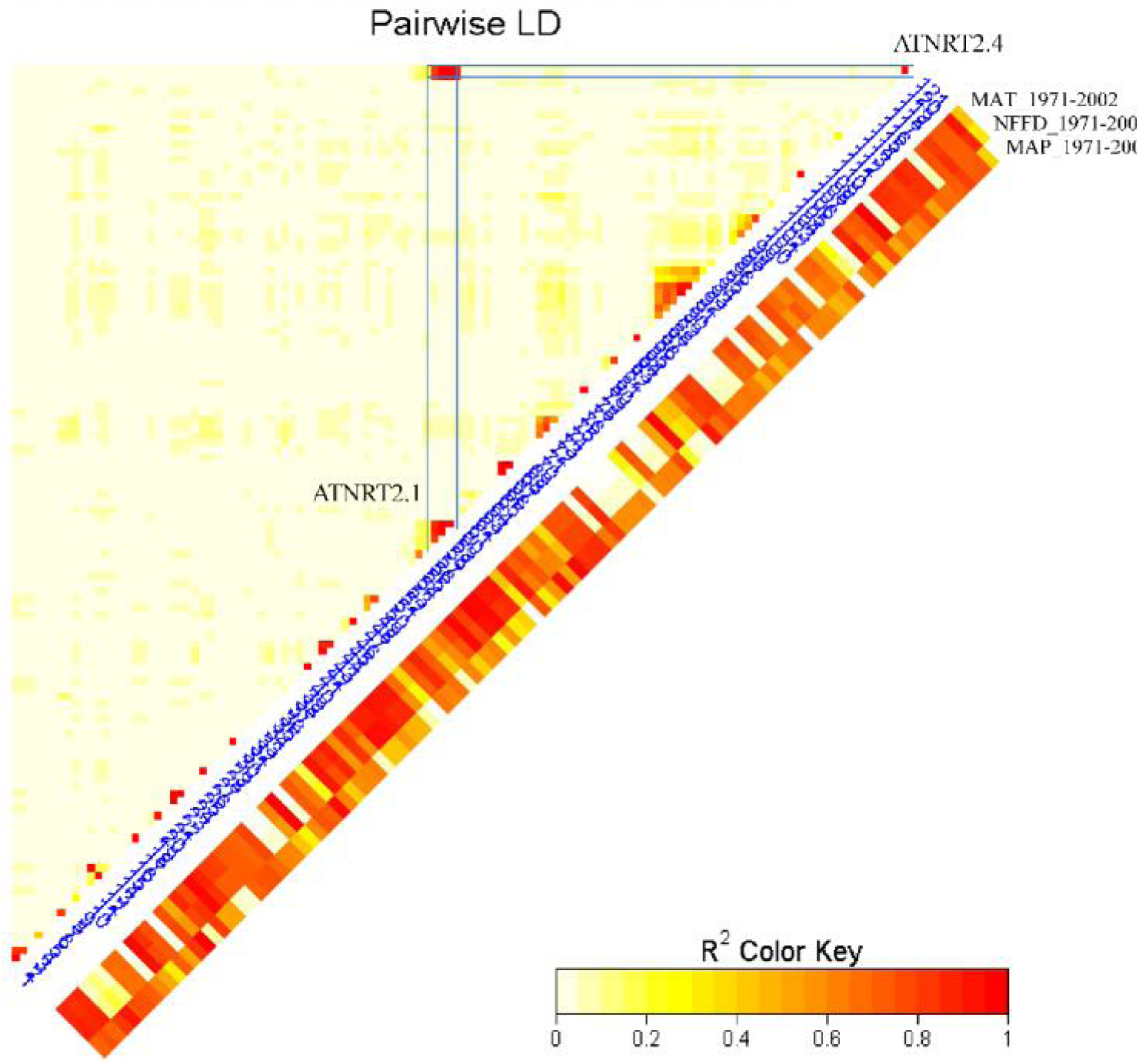
Linkage disequilibrium between 121 identified *F*_*ST*_ outlier loci and relationship between *F*_*ST*_ outlier allele frequencies and climate variables in *P. trichocarpa*. Simple linear regression (R^2^) of allelic frequencies (following arcsine transformation) on temperature and precipitation, respectively (mean annual temperature in °C: MAT_1971-2002; number of frost-free days: NFFD_1971-2002 and mean annual precipitation in mm: MAP_1971-2002, observed between yrs 1971-2002) calculated among the four distinct climate clusters (Fig. 1); Note: POPTR_0143s00200 was recently re-annotated to Potri.009G008500 and both genes are now assembled on chromosome 9 within 50kb of each other (new poplar genome assembly Phytozyme v3). Both sequences are now described as tandem gene pair *PTNRT2.4A* (alias Potri.009G008600) and *PTNRT2.4B* (alias Potri.009G008500) with 97% DNA sequence similarity (Bai *et al*., 2013).

The order of loci follows:

1. scaffold_1_27485620
2. scaffold_1_27487874
3. scaffold_1_27488119
4. scaffold_1_33628533
5. scaffold_1_33632379
6. scaffold_1_37065304
7. scaffold_1_37410840
8. scaffold_1_37410856
9. scaffold_1_45757179
10. scaffold_1_45758739
11. scaffold_2_127966
12. scaffold_2_128416
13. scaffold_2_128432
14. scaffold_2_130506
15. scaffold_2_10949533
16. scaffold_2_13035475
17. scaffold_3_14135487
18. scaffold_3_14135542
19. scaffold_3_19339785
20. scaffold_3_19747482
21. scaffold_3_19750521
22. scaffold_4_17161026
23. scaffold_4_17161413
24. scaffold_4_17162655
25. scaffold_5_88127
26. scaffold_5_12339685
27. scaffold_5_12344723
28. scaffold_5_16487025
29. scaffold_5_16811923
30. scaffold_5_19211088
31. scaffold_5_19211834
32. scaffold_5_19953723
33. scaffold_5_22633044
34. scaffold_6_2485373
35. scaffold_6_2489698
36. scaffold_6_3249232
37. scaffold_6_6390362
38. scaffold_6_6436509
39. scaffold_6_23299767
40. scaffold_6_24631540
41. scaffold_6_24634215
42. scaffold_6_25893186
43. scaffold_6_25893407
44. scaffold_6_25893900
45. scaffold_7_74879
46. scaffold_7_178643
47. scaffold_7_179188
48. scaffold_7_808919
49. scaffold_7_809632
50. scaffold_7_811143
51. scaffold_8_805284
52. scaffold_8_6567373
53. scaffold_8_9267412
54. scaffold_9_1379696
55. scaffold_9_1599746
56. scaffold_9_1606213
57. scaffold_9_1676227
58. scaffold_9_1676590
59. scaffold_9_1678624
60. scaffold_9_1678826
61. scaffold_9_2160922
62. scaffold_9_2563600
63. scaffold_9_2677917
64. scaffold_9_2679340
65. scaffold_9_2687811
66. scaffold_9_3795784
67. scaffold_9_3798176
68. scaffold_9_3800384
69. scaffold_10_255159
70. scaffold_10_20168770
71. scaffold_10_21246081
72. scaffold_10_21249991
73. scaffold_10_21253673
74. scaffold_10_21451968
75. scaffold_11_145058
76. scaffold_11_295988
77. scaffold_11_15084939
78. scaffold_11_15084942
79. scaffold_11_18477497
80. scaffold_12_1811250
81. scaffold_12_1811719
82. scaffold_12_1812031
83. scaffold_13_14296993
84. scaffold_14_12173467
85. scaffold_14_12173560
86. scaffold_14_12927245
87. scaffold_15_133408
88. scaffold_15_247054
89. scaffold_15_247527
90. scaffold_15_247811
91. scaffold_15_267849
92. scaffold_15_268612
93. scaffold_15_342410
94. scaffold_15_382827
95. scaffold_15_512479
96. scaffold_15_630677
97. scaffold_15_703349
98. scaffold_15_704562
99. scaffold_15_718240
100. scaffold_15_719540
101. scaffold_15_719682
102. scaffold_15_910808
103. scaffold_15_1006871
104. scaffold_15_13596400
105. scaffold_15_13618770
106. scaffold_15_13808656
107. scaffold_15_13808709
108. scaffold_15_13889772
109. scaffold_17_724384
110. scaffold_17_5220579
111. scaffold_17_12392905
112. scaffold_17_12436896
113. scaffold_18_1110947
114. scaffold_18_2565040
115. scaffold_19_5985766
116. scaffold_19_12221032
117. scaffold_19_12484019
118. scaffold_19_15299925
119. scaffold_21_280997
120. scaffold_143_2955
121. scaffold_143_3026

